# Unraveling the Evolutionary Trajectory of LHCI in Red-Lineage Algae: Conservation, Diversification, and Neolocalization

**DOI:** 10.1101/2024.03.16.585336

**Authors:** Minoru Kumazawa, Kentaro Ifuku

**Author notes:** Corresponding Author: Kentaro Ifuku.

## Abstract

Red algae and the secondary symbiotic algae that engulfed a red alga as an endosymbiont are called red-lineage algae. They comprise key marine taxa including diatoms, Haptophyta, and Cryptophyta. Several photosystem (PS) I–light-harvesting complex I (LHCI) structures have been reported from red-lineage algae —two red algae *Cyanidioschyzon merolae* (Cyanidiophyceae), *Porphyridium purpureum* (Rhodophytina), a diatom *Chaetoceros gracilis* and a Cryptophyte *Chroomonas placoidea*. Here, we clarified the orthologous relation of LHCIs in red-lineage algae by combining a detailed phylogenetic analysis of LHCIs and the structural information of PSI–LHCI. We found that the seven Lhcr groups in LHCI are conserved in Rhodophytina; Furthermore, during both genome reduction in Cyanidioschyzonales of red algae and endosymbiosis leading to Cryptophyta, some LHCIs were lost and replaced by existing or differentiated LHCIs. Especially in Cryptophyta, uniquely diversified Lhcrs form three sets of heterotrimers contributed to the expansion of the antenna size of PSI, supporting the modern ecological success of this taxon. We denominated “neolocalization” to these examples of flexible reorganization of LHCIs. This study provides new insights into the evolutionary process of LHCIs associated with PSI in the red-lineage algae and clarifies the need for both molecular phylogeny and structural information to elucidate the plausible evolutionary history of LHCI.

## Introduction

Oxygenic photosynthetic organisms, such as cyanobacteria, algae, and terrestrial plants, play an essential role in capturing sunlight, producing organic matter, and maintaining life on Earth both underwater and on land. Among the various eukaryotic photosynthetic organisms, algae and terrestrial plants acquired chloroplasts through endosymbiosis with cyanobacteria (Delwiche, 1999). These photosynthetic organisms possessing primary plastids form Archaeplastida, and are divided into three groups, Rhodophyta (red algae), Viridiplantae (green algae and terrestrial plants), and Glaucophyta. Several secondary or tertial endosymbiotic events led to diversified eukaryotic algae. For instance, red-lineage secondary endosymbiotic algae acquired plastids derived from red algae and include key marine taxa including diatoms and Haptophytes, which dominate in modern oceans (Pierella Karlusich *et al*., 2020).

To enable more efficient light capture, photosynthetic organisms possess peripheral light-harvesting antennas around the two photosystems. In eukaryotic photosynthetic organisms, these antennas are protein complexes holding light-harvesting pigments, which transfer the excitation energy acquired from the light to the photosystems through excitation energy transfer (Croce and van Amerongen, 2020). Red algae have a phycobilisome, a superficial light-harvesting antenna complex on the stromal side of photosystem (PS) II, and two-dimensionally coordinated transmembrane light-harvesting pigment-protein complexes (LHCs) associated with PSI (Pi *et al*., 2018; You *et al*., 2023; Wolfe *et al*., 1994; Marquardt and Rhiel, 1997). LHCs bind various types of chlorophylls and carotenoids as light-harvesting pigments and serve as light-harvesting antennas in red and green algae, land plants, red-lineage secondary endosymbiotic algae, green lineage secondary endosymbiotic algae, and dinoflagellates (Koziol *et al*., 2007; Büchel, 2015).

Red algal LHCs contain chlorophyll *a* and zeaxanthin as a carotenoid, while most LHCs of red-lineage secondary endosymbiotic algae include chlorophyll *a* and *c*; carotenoids depend on the taxonomic group. In fact, LHCs are named after their binding carotenoids (Büchel, 2015). For example, diatoms and Haptophytes contain fucoxanthin or 19’-hexanoyloxy fucoxanthin as major carotenoid in their LHCs, thus their LHCs are called fucoxanthin chlorophyll *a*/*c*-binding proteins (FCPs). Among red-lineage secondary endosymbiotic algae, diatoms utilize FCPs as peripheral antennas for both PS I and II (Nagao *et al*., 2020; Nagao *et al*., 2019; Nagao *et al*., 2022; Xu *et al*., 2020; Wang *et al*., 2019). Similar light-harvesting systems probably exist in other Stramenopiles and Haptophytes. At least, Eustigmatophyceae, belonging to Stramenopiles, utilize LHCs for light-harvesting for both PS (Umetani *et al*., 2018).

Based on molecular phylogeny, the LHCs of red-lineage algae are divided into six subfamilies: Lhcr, Lhcz, Lhcq, Lhcf, Lhcx, and CgLhcr9 homologs (Kumazawa *et al*., 2022). Some Stramenopiles, including Eustigmatophyceae and Phaeophyceae, as well as Chromera from Alveolate have another LHC subfamily called red-shifted *Chromera* light-harvesting proteins (Red-CLH) (Bína *et al*., 2014; Umetani *et al*., 2018). Red algae only have the Lhcr subfamily, Stramenopiles and Haptophytes possess all six subfamilies of the red-lineage LHCs, while Cryptophytes only have Lhcr and Lhcz subfamilies.

Recently, the advancement of cryoelectron microscopy structural analysis allowed discerning the structures of the PS–peripheral light-harvesting antenna supercomplexes of red-lineage algae. In red algae, the *Cyanidioschyzon merolae* PSI–LHCI supercomplex (Pi *et al*., 2018) and the *Porphyridium purpureum* phycobilisome–PSII–PSI–LHCI megacomplex (You *et al*., 2023) have been reported. In red-lineage secondary endosymbiotic algae, molecular-level structural PS models have been reported in diatoms and a Cryptophyte. For instance, the diatom *Chaetoceros gracilis* PSI–FCPI supercomplexes (Nagao *et al*., 2020; Xu *et al*., 2020) and *C. gracilis* PSII–FCPII supercomplexes (Nagao *et al*., 2019; Wang *et al*., 2019; Nagao *et al*., 2022) have been reported. In addition, PSII–FCPII structures from centric diatoms *Thalassiosira pseudonana* and *Cyclotella meneghiniana* were recently reported (S., Zhao *et al*., 2023; Feng *et al*., 2023). The LHCs of Cryptophytes are called alloxanthin-chlorophyll *a*/*c*-binding proteins (ACPs); the structure of the *Chroomonas placoidea* PSI– ACPI has been recently reported (Zhao et al., 2023).

With the identification of LHCs in the PSI–LHCI structural models recently reported, it is now possible to evaluate the evolutionary process of the molecular assembly of LHCI associated with PSI. The molecular assembly model of the red-lineage PSI–LHCI has been discussed only based on spatial arrangements of the subunits present in the structures (Bai *et al*., 2021; L.,-S., Zhao *et al*., 2023). However, an evolutionary model of the photosynthetic supercomplex should comprise both molecular phylogeny and structural information. Such an integrative understanding of the complex structures and molecular phylogenies of primitive species has been attempted in the green lineage (Neilson and Durnford, 2010). Loss and gain of LHC subfamilies during the evolutionary history of the red-lineage algae were investigated through phylogenetic analysis of diatom LHCs (Kumazawa et al. 2022). A red-lineage chlorophyll *a*/*b*-binding-like protein (RedCAP), a distinctive family of the LHC superfamily, is conserved in PSI–LHCI of Rhodophytina red algae, Cryptophytes, and diatoms (Engelken *et al*., 2010; Sturm *et al*., 2013; Xu *et al*., 2020; You *et al*., 2023; L.,-S., Zhao *et al*., 2023). However, the evolutionary model of the red algal LHCI and RedCAP remains incomplete because of the limited structural information, insufficient genome and transcriptome information (until recently), and lack of detailed molecular phylogeny at the ortholog level (Hoffman *et al*., 2011; Dittami *et al*., 2010).

In this study, we performed a molecular phylogenetic analysis to clarify the orthology of LHCIs in red-lineage algae using recently reported genomes and transcriptomic data. The detailed molecular phylogeny of the red-lineage LHCI, specifically Lhcrs, is combined with new PSI–LHCI structural models from red and red-lineage algae. This has uncovered conservation, diversification, and differentiation of the molecular assembly of LHCIs, especially in red algae and Cryptophytes. Based on our analyses, we propose a new evolutionary trajectory of LHCI proteins associated with PSI in red-lineage algae.

## Results

### Molecular Phylogeny of Red Algal LHCI

Red algae possess two types of membrane-spanning light-harvesting pigment-protein complexes in PSI; LHCs that belong to the Lhcr subfamily and a RedCAP, part of the LHC superfamily (Engelken *et al*., 2010; Sturm *et al*., 2013; You *et al*., 2023). In contrast, in PSII they have a large membrane-peripheral light-harvesting protein supercomplex, known as phycobilisome. To elucidate the molecular phylogeny of the LHC family in PSI, Lhcr sequences from a broad lineage of red algae were collected minimizing as much as possible any taxonomic biases.

Ancient Rhodophytina is the original endosymbiont of red-lineage secondary endosymbiotic algae (Fig. 1) (Yoon *et al*., 2002; Kim *et al*., 2017). After secondary endosymbiosis, the earliest divergent event divided red algae into two major groups: Rhodophytina and Cyanidiophyceae (or Cyanidiophytina) (Yang *et al*., 2016; Park *et al*., 2023): The former includes classes such as Porphyridiophyceae, Stylonematophyceae, and Compsopogonophyceae with the subclades of Rhodellophyceae, Bangiophyceae, and Florideophyceae (Yang *et al*., 2016; Borg *et al*., 2023). The latter group, Cyanidiophyceae, contains the orders Galdieriales, Cavernulicolales, Cyanidiales, and Cyanidioschyzonales (Park *et al*., 2023). Among them, Galdieriales is considered as the earliest diverged order. Genomes or transcriptomes are available for all orders except Cavernulicolales; further, LHC sequences could be obtained through homology searches. Additionally, we obtained the sequences of putative LHCI (Lhca) associated with PSI in *Prasinoderma coloniale*—a member of the Prasinodermophyta class representing the earliest divergence within the primary green lineage— (Li *et al*., 2020).

**Figure 1.**
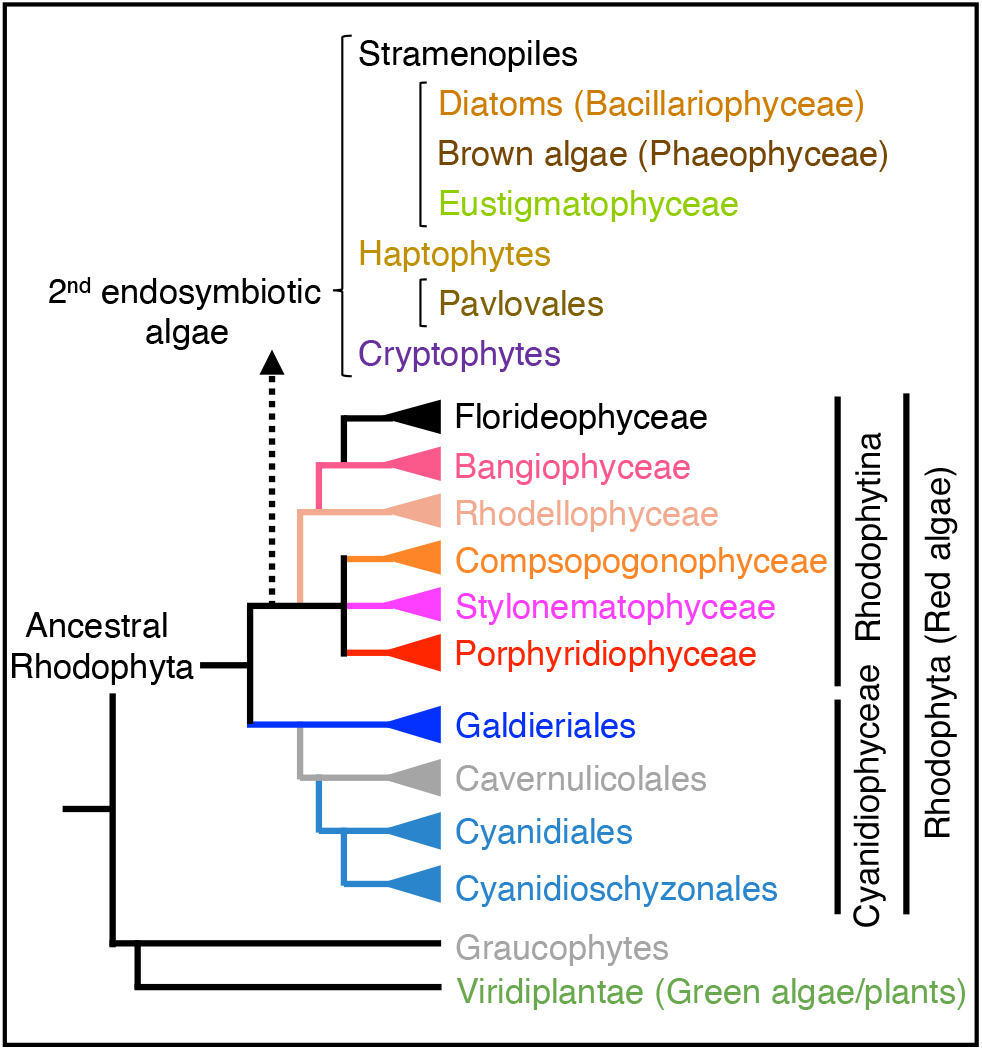
Simplified phylogeny of Rhodophyta (red algae) and classification of red-linage secondary endosymbiotic algae. The phylogeny of Rhodophyta is based on Yang et al. (2016) and Park et al. (2023). The secondary endosymbiosis node in the tree is based on Yoon et al. (2002) and Kim et al. (2017). The root of the tree is the ancestor of Archaeplastida. “Ancestral Rhodophyta” in the figure represents the last common ancestor of extant Rhodophyta.

Next, a molecular phylogenetic tree was constructed using the obtained Lhcr sequences from red algae and Lhca sequences from the green alga *P. coloniale* (Fig. 2). There are seven groups of Lhcrs (group I–VII) from red algae belonging to Rhodophytina, including *Porphyridium purpureum*. Each group contains one *P. purpureum* Lhcr; Group VII, III, and VI containing PpLhcr3, PpLhcr4, and PpLhcr6 show monophyly, and Group III and VI are sister groups to group VII. However, the distribution of Cyanidiophyceae Lhcrs is different: Galdieriales, belonging to Cyanidiophyceae, has Lhcrs classified into six including group I and IV–VII in addition to the Galdieriales-specific clade. This unique clade is a sister clade to group I. Galdieriales lacks group II and -III LHCs. Cyanidiales and Cyanidioschyzonales have only three orthologs to those of Rhodophytina, and these Lhcrs –*Cyanidioschyzon merolae* Lhcr1 (hereafter CmLhcr1), CmLhcr2, and CmLhcr3– which belong to group V, VI and VII, respectively.

**Figure 2.**
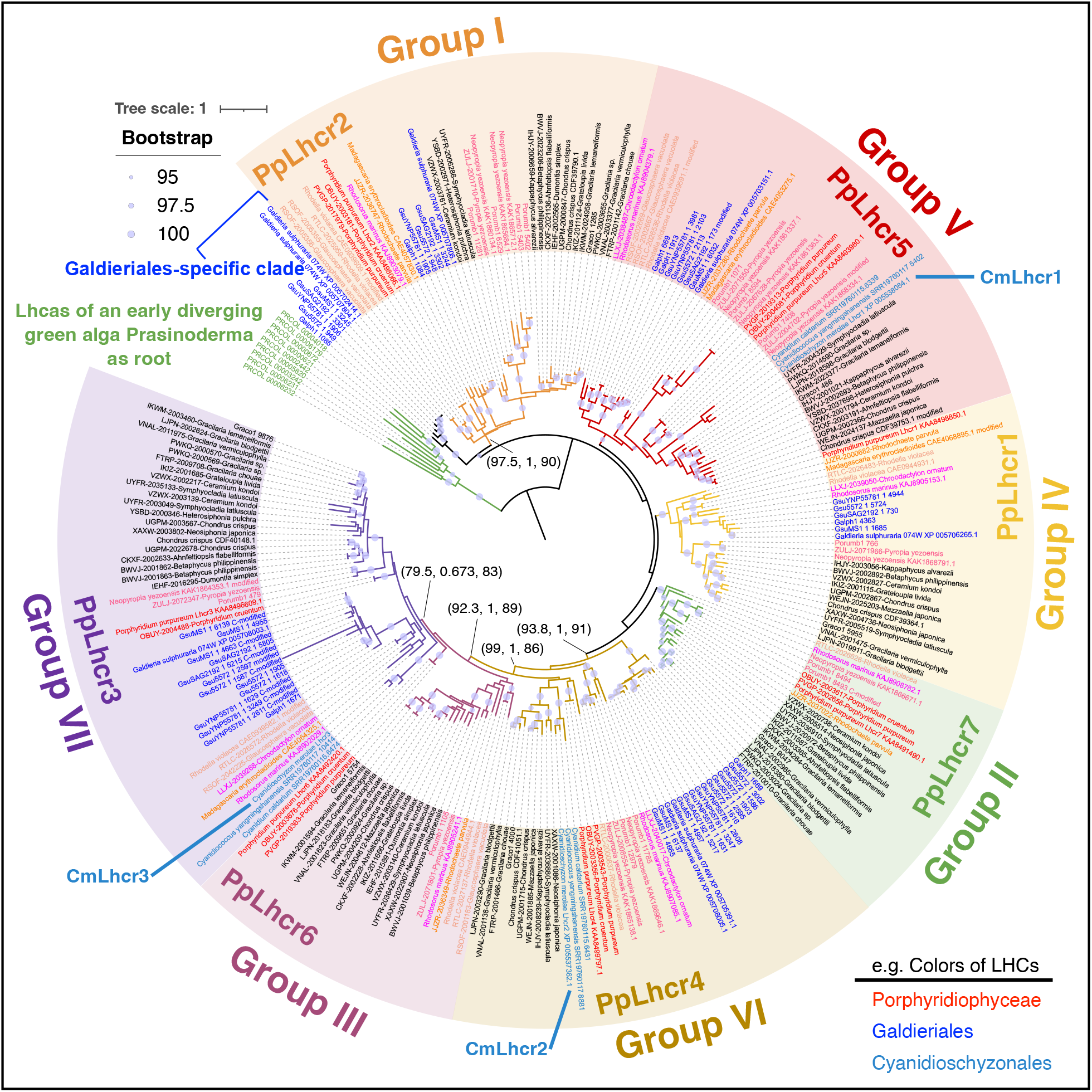
Molecular phylogenetic tree of Lhcr of red algae. The phylogenetic tree was constructed using IQ-TREE2 and rooted at the outgroup of Lhcas of an early diverging green alga, *Prasinoderma coloniale*. A total of 261 sequences with 341 amino-acid sites were used, and the Q.pfam+F+I+R7 model was selected according to Bayesian information criterion scores. Circles on the node indicate ultrafast bootstrap support (≥95%). Numbers in parentheses are SH-aLRT support (%) / aBayes support / ultrafast bootstrap support (%). LHC colors correspond to the taxonomy in Figure 1. Short names of molecular species are described in Supplemental Table 1. For example, *C. gracilis* Lhcr1 is shown as CgLhcr1.

### PSI–LHCI in “Primitive Red Algae” Cyanidioschyzonales

In a previous structural model of PSI–LHCI for *Cyanidioschyzon merolae* belonging to Cyanidioschyzonales (Pi *et al*., 2018), there were five LHCs attached to PSI, including two copies of CmLhcr1 (r1 and r1*) and CmLhcr2 (r2 and r2*) in addition to one CmLhcr3 (r3) (Fig. 3). The structure of the phycobilisome–PSII–PSI–LHCI megacomplex (Fig. 3) has been reported in another red alga, *P. purpureum* belonging to Porphyridiophyceae (You *et al*., 2023); it has eight LHCIs (PpLhcr1–7 and RedCAP) around PSI.

**Figure 3.**
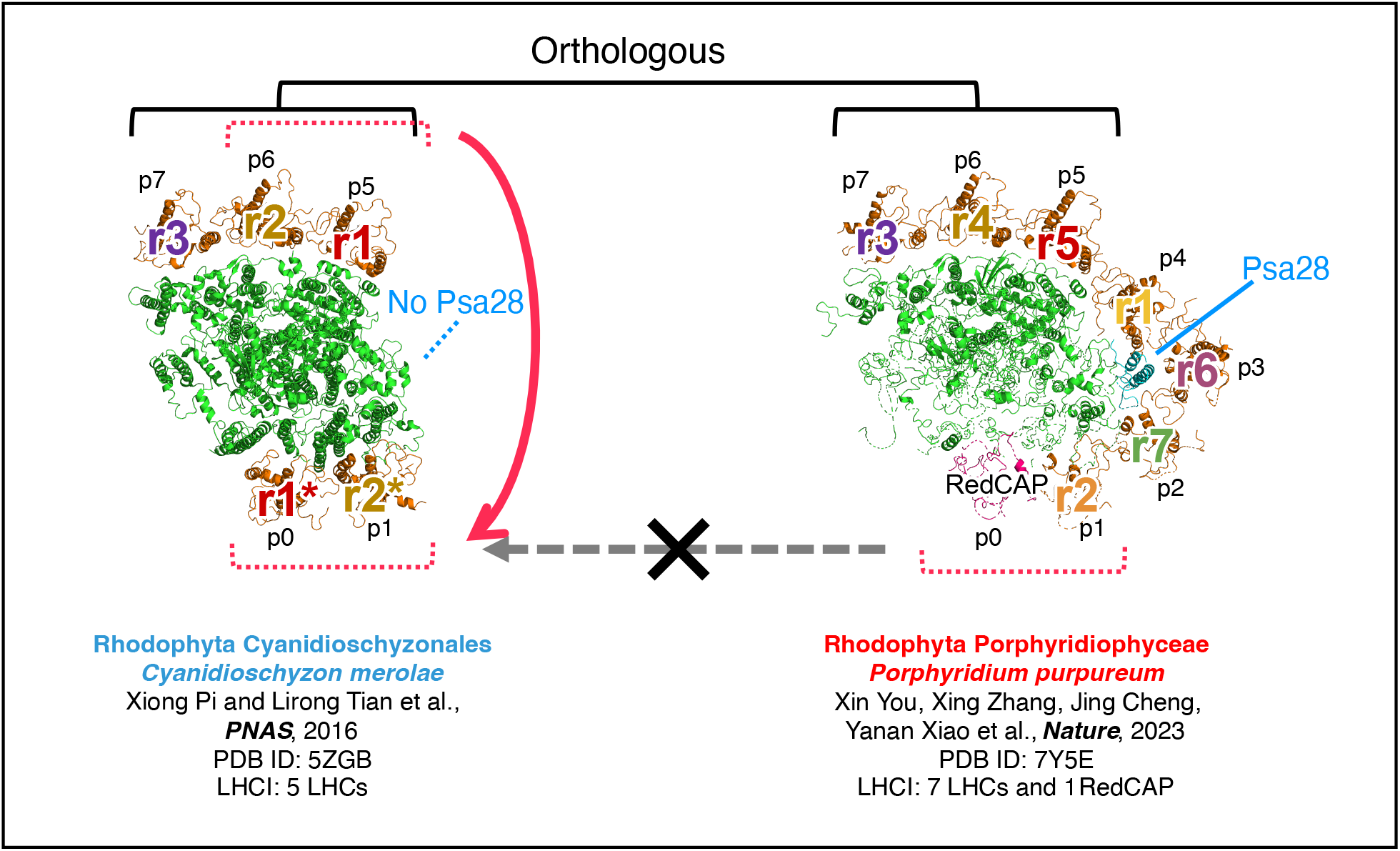
Estimated scheme of PSI-LHCI evolution in *Cyanidioschyzon.* The left PSI–LHCI structure corresponds to the Cyanidioschyzonales *Cyanidioschyzon merolae* and the right PSI–LHCI to the Porphyridiophyceae *Porphyridium purpureum*. The names of LHCIs are adapted from the original papers (Xiong Pi and Lirong Tian et al., 2016; Xin You, Xing Zhang, Jing Cheng, Yanan Xiao et al., 2023) and colored according to Figure 2.

From the stromal side, in the *P. purpureum* PSI–LHCI part, positions of eight molecular LHCI species (r1–r7, and RedCAP) are labeled counterclockwise as positions 0–7 (p0–p7) (Fig. 3). Accordingly, the positions of LHCI in *C. merolae* is labeled as p0, p1, and p5–p7 for two CmLhcr1 (r1 and r1*), two CmLhcr2 (r2 and r2*), and CmLhcr3 (r3) (Pi *et al*., 2018). *C. merolae* PSI–LHCI does not have an LHC at positions p2–p4. The loss of Psa28 (also called PsaR) in the genome of *C. merolae* implies that Psa28 would be crucial for LHCI binding at p2–p4 (You *et al*., 2023).

The Lhcrs at p5–p7 of both species seem to be conserved from their common ancestor. Our phylogenetic analysis suggests that CmLhcr1–3 at p5–p7 in *C. merolae* PSI–LHCI belong to groups V, VI, and VII, respectively, and that PpLhcr5, PpLhcr4, and PpLhcr3 at p5–p7 in *P. purpureum* PSI–LHCI belong to group V, VI and VII, which are orthologs to CmLhcr1–3 (Fig. 2 and 3). In contrast, significant changes are observed between the two species in Lhcrs at p0 and p1 (Fig. 3): *P. purpureum* PSI binds RedCAP at p0, while that of *C. merolae* binds CmLhcr1 at p0 in addition to p5; *P. purpureum* binds PpLhcr2 (group I Lhcr) at p1, while *C. merolae* binds CmLhcr2 (group VI Lhcr) at p1 as well as p6. Importantly, PpLhcr2 and CmLhcr2 are not orthologs in the phylogenetic tree, although they bind at the same position; instead, CmLhcr2 and PpLhcr4 at p6 belong to group VI and they show an orthologous relationship in the tree. Given that RedCAP is conserved in Galdieriales which is an early diverged taxa of Cyanidiophyceae, RedCAP is lost in *C. merolae* (Engelken *et al*., 2010; Sturm *et al*., 2013). *C. merolae* lost RedCAP, group I–IV LHCIs, and a related PSI subunit, and complemented positions p0 and p1 with group V and VI LHCIs to, at least partially, maintain the antenna size for PSI.

### Molecular Phylogeny of Lhcrs in Red-Lineage Algae

Cryptophytes do not possess the Lhcf subfamily or phycobilisome, but they do contain phycobiliproteins in the thylakoid lumen and LHCs in the thylakoid membrane (Spear-Bernstein and Miller, 1989). Cryptophyte LHC is called chlorophyll *a*/*c* (CAC) proteins or alloxanthin-chlorophyll *a*/*c* proteins (ACPs) named after their pigments and bound to both PSI and PSII (Kereïche *et al*., 2008; Kaňa *et al*., 2012; L.,-S., Zhao *et al*., 2023). The PSI–LHCI and PSII–LHCII of diatoms and Haptophytes are called PSI–FCPI and PSII–FCPII, respectively, because of the bound pigments in LHCs. The classification of LHCs in red-lineage secondary endosymbiotic algae was described in our previous research (Kumazawa *et al*., 2022). The diatom FCPI comprises Lhcrs as well as the Lhcqs, CgLhcr9, which is distinct from the Lhcr subfamily, presumably some Lhcfs, and a RedCAP (Nagao *et al*., 2020; Xu *et al*., 2020; Kumazawa *et al*., 2022; Calvaruso *et al*., 2020). Unlike red algal PSII with phycobilisome, diatoms predominantly have the Lhcfs located around PSII, and one unique Lhcr (CgLhcr17 homolog) closely associated with the PSII core (Nagao *et al*., 2019; Nagao *et al*., 2022; Wang *et al*., 2019; Kumazawa *et al*., 2022; Calvaruso *et al*., 2020). Molecular phylogenetic analysis suggests that Haptophytes also possess LHC subfamily compositions similar to diatoms, implying that they may have analogous PSI–FCPI and PSII–FCPII (Kumazawa *et al*., 2022). To elucidate how the Lhcrs in these algae were generated from red algae during secondary endosymbiosis, LHC sequences were obtained from a wide variety of Cryptophytes, Stramenopiles, and Haptophytes, and we performed molecular phylogenetic analysis on the obtained Lhcr sequences using *Prasinoderma* Lhcas as root of the tree (Fig. 4).

**Figure 4.**
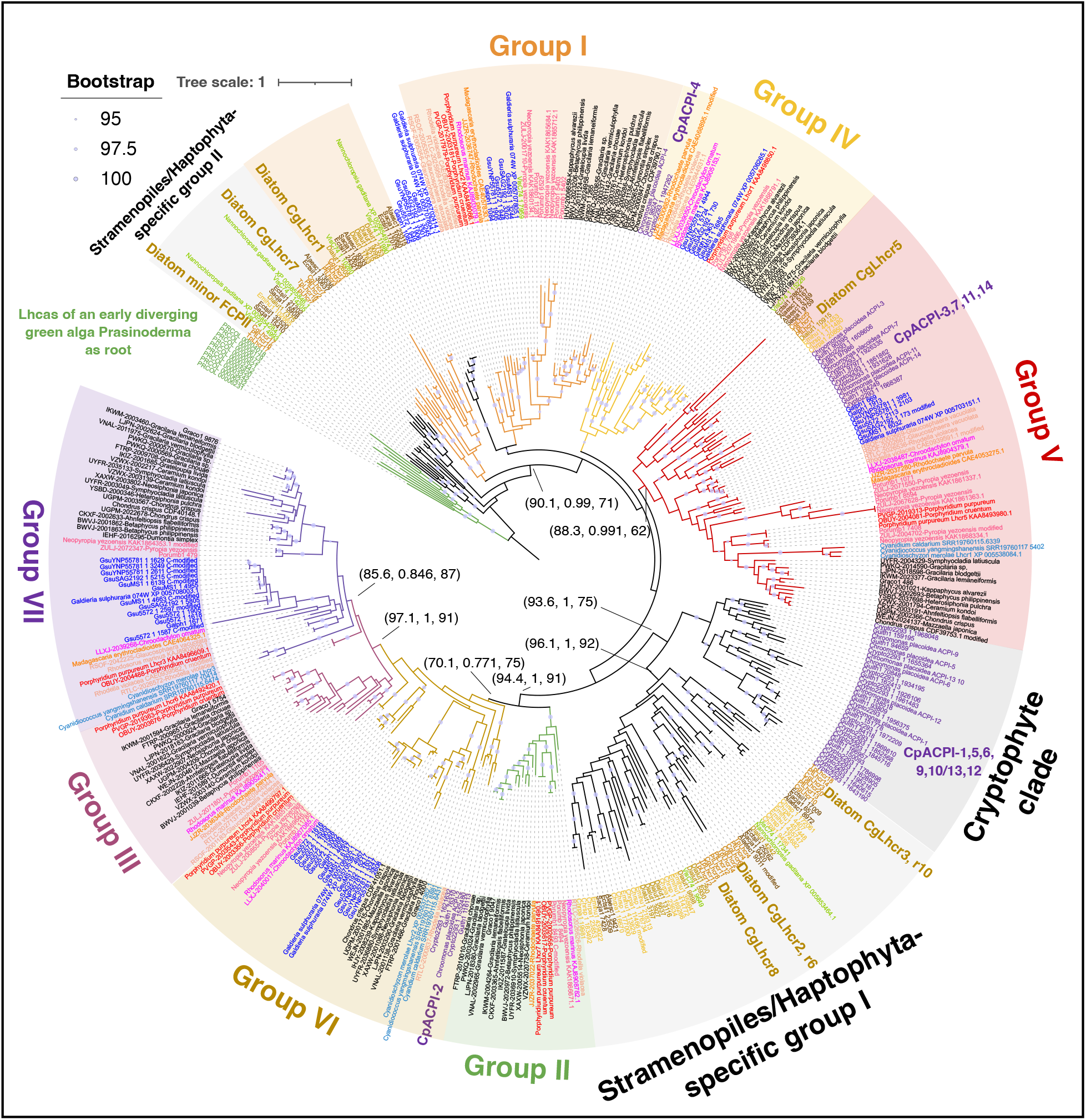
Molecular phylogenetic tree of Lhcr of red-lineage algae. The phylogenetic tree was inferred using IQ-TREE2 and rooted at the outgroup of Lhcas from an early diverging green alga, Prasinoderma coloniale. A total of 418 sequences with 489 amino-acid sites were used, and the Q.pfam+F+R7 model was selected according to Bayesian information criterion scores. Circles on the node indicate ultrafast bootstrap support (≥95%). Numbers in parentheses are SH-aLRT support (%) / aBayes support / ultrafast bootstrap support (%). LHC colors correspond to the taxonomy in Figure 1. Short names of molecular species are described in Supplemental Table 1.

LHCIs of Cryptophytes and Stramenopiles show different conservation patterns in the phylogenetic tree, suggesting different evolutionary processes of LHCIs (Fig. 4). The phylogenetic tree of red-lineage LHCIs includes group I–VII in addition to one Cryptophyte-specific clade and two Stramenopiles/Haptophyta-specific clades. Cryptophytes, including *Chroomonas placoidea*, have ACPIs in groups IV–VI in addition to the Cryptophyte-specific clade. Group IV and VI contain CpACPI-4 and CpACPI-2, respectively. Group V contains not only CpACPI-3 but also CpACPI-7, 11, and 14. Groups I–III and VII are absent in Cryptophytes. The Cryptophyte-specific clade contains CpACPI-1, 5, 6, 9, 10/13, and 12; ACPI-6 and ACPI-10/13 are closely related and form one group, whereas CpACPI-1, 5, 9 forms the other.

Stramenopiles, including the diatom *Chaetoceros gracilis*, have CgLhcr1 in group I and CgLhcr5 in group V in addition to LHCIs in two Stramenopiles/Haptophyta-specific groups. The clade including CgLhcr1 is a sister clade of the group I clade of red algae, suggesting that this clade can be included in group I. One Stramenopiles/Haptophyta-specific group contains CgLhcr2, r3, r6, r8, and r10, while the other includes CgLhcr7 and r17. CgLhcr17 is a monomeric FCPII directly associated with PSII (Kumazawa *et al*., 2022; Nagao *et al*., 2022). The Lhcrs of other Stramenopiles showed the same distribution pattern in the groups as that of diatom Lhcrs. Group I includes Pavlovales Lhcrs from Haptophytes, whereas Stramenopiles/Haptophyta-specific groups possess Lhcrs from other taxa of Haptophyta. Altogether, Lhcrs of red-lineage secondary endosymbiotic algae are present not only in red algal Lhcr groups but also in unique clades in the phylogenetic tree.

### Cryptophyte PSI–ACPI and diatom PSI–FCPI showing LHCI rearrangement

Conservation and replacement of LHCIs (ACPIs) from ancestral red algae can be estimated when we combined the structure of Cryptophyte *Chroomonas placoidea* PSI– ACPI with the phylogenetic analysis. *C. placoidea* PSI–ACPI contains one RedCAP molecule and 13 Lhcrs (Fig. 5) (Zhao et al., 2023). It also has an unknown subunit and ACPI-S, which appears to interconnect ACPIs and is not found in red algal PSI–LHCI and diatom PSI–FCPI. From the stromal side, the binding positions of LHCI (ACPIs) can be assigned as p0–p8 similar to the red algal PSI–LHCI (Fig. 3). RedCAP locates at p0 as observed in *P. purpureum*. ACPI-4, 3, and 2 at p4, p5, and p6 belong to groups IV, V, and VI, respectively, as did LHCIs in *P. purpureum* PSI–LHCI. In contrast, ACPI-7 at p1 belongs to group V in *C. placoidea,* while p1 is occupied by group I PpLhcr2 in *P. purpureum* PSI–LHCI. Furthermore, p2, p3, and p7 are occupied by ACPI-6, -5, and -1 belonging to the Cryptophyte-specific clade.

**Figure 5.**
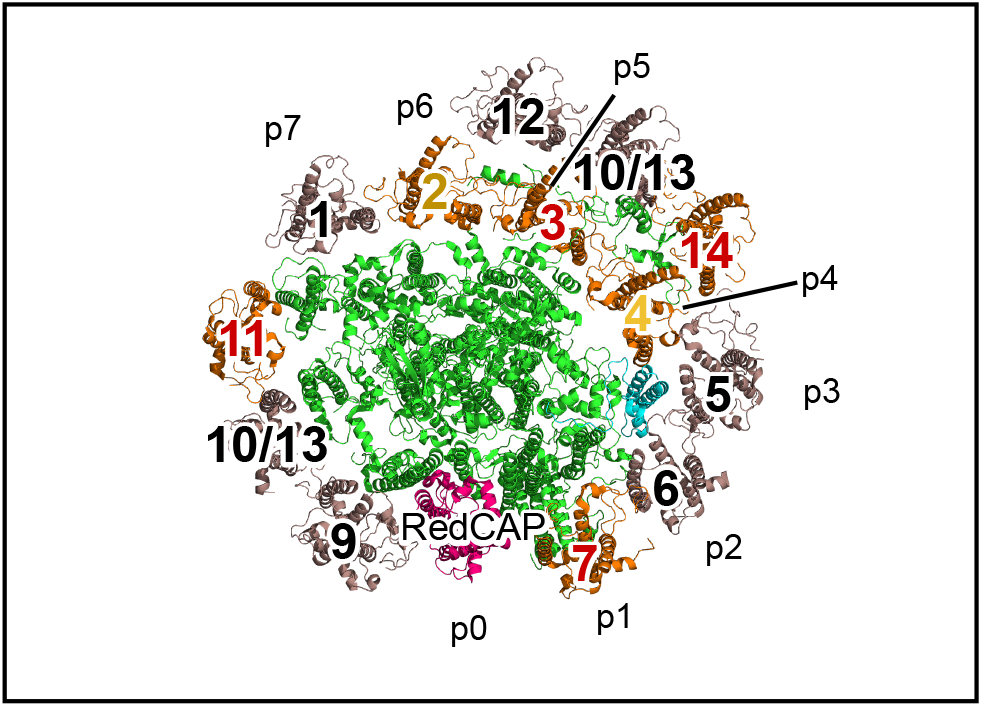
PSI–LHCI structures of Cryptophyte *Chroomonas placoidea* PSI–ACPI supercomplex. Top view of the supercomplex from the stromal side. The number on the structure corresponds to that of CpACPI proteins. The colors of ACPI numbers correspond to the groups in Figure 4. PSI core subunits except Psa28 (alternatively known as PsaR) are shown in green and Psa28 is cyan, respectively. RedCAP protein is shown in red and Lhcr proteins in LHCI orthologous/homologous to the red algal Lhcrs are colored orange. Lhcrs belonging to the Cryptophyte-specific clade are shown in brown.

Interestingly, *Chroomonas placoidea* PSI–ACPI has three sets of adjacent three ACPIs (heterotrimer): ACPI-7–6–5, ACPI-11–10/13–9, and 14–10/13–12 (Fig. 5). ACPI-7, 11, and 14 in each heterotrimer belong to group V. ACPI-3 belonging to group-V ACPIs would be rather ancestral LHCI because it binds at p3, meaning a true ortholog of group-V LHCI in red algae (ex. PpLhcr5). ACPI-7, 11, and 14 belonging to group V should be derivatives of ancestral ACPI-3. Other two ACPIs in each heterotrimer belong to the unique Cryptophyte-specific clade, wherein *C. placoidea* ACPIs can be divided into two groups by focusing on monophyly: ACPI-6, 10/13, and ACPI-5, 9, 12 (Fig. 4). Thus, three sets of heterotrimers comprise three ACPIs from the respective groups, which would result from gene duplication.

According to the above analyses, the following model can be proposed for the establishment of Cryptophyte PSI–ACPI (Fig. 6): After losing many Lhcr genes during secondary endosymbiosis of red algae, Cryptophytes only preserved three molecular species of Lhcr (ACPI-2, ACPI-3, and ACPI-4) and a RedCAP; at this point, the group V LHCI—an ancestral ACPI-3 at p5—diversified forming a Cryptophyte-specific clade in the Lhcr subfamily: one Cryptophyte-specific Lhcrs binds at p7 as ACPI-1; three sets of heterotrimers of ACPIs, including one group V Lhcr and two Cryptophyte-specific Lhcrs, bind to restore the antenna size of PSI. This molecular evolutionary model of *C. placoidea* PSI–ACPI (Fig. 6) contradicts the current model of PSI–LHCI complex evolution in the red-lineage, solely based on LHCI compositions (Zhao et al., 2023). That is, Cryptophyte PSI–ACPI is not an evolutional intermediate between red algal PSI–LHCI and diatom PSI–FCPI.

**Figure 6.**
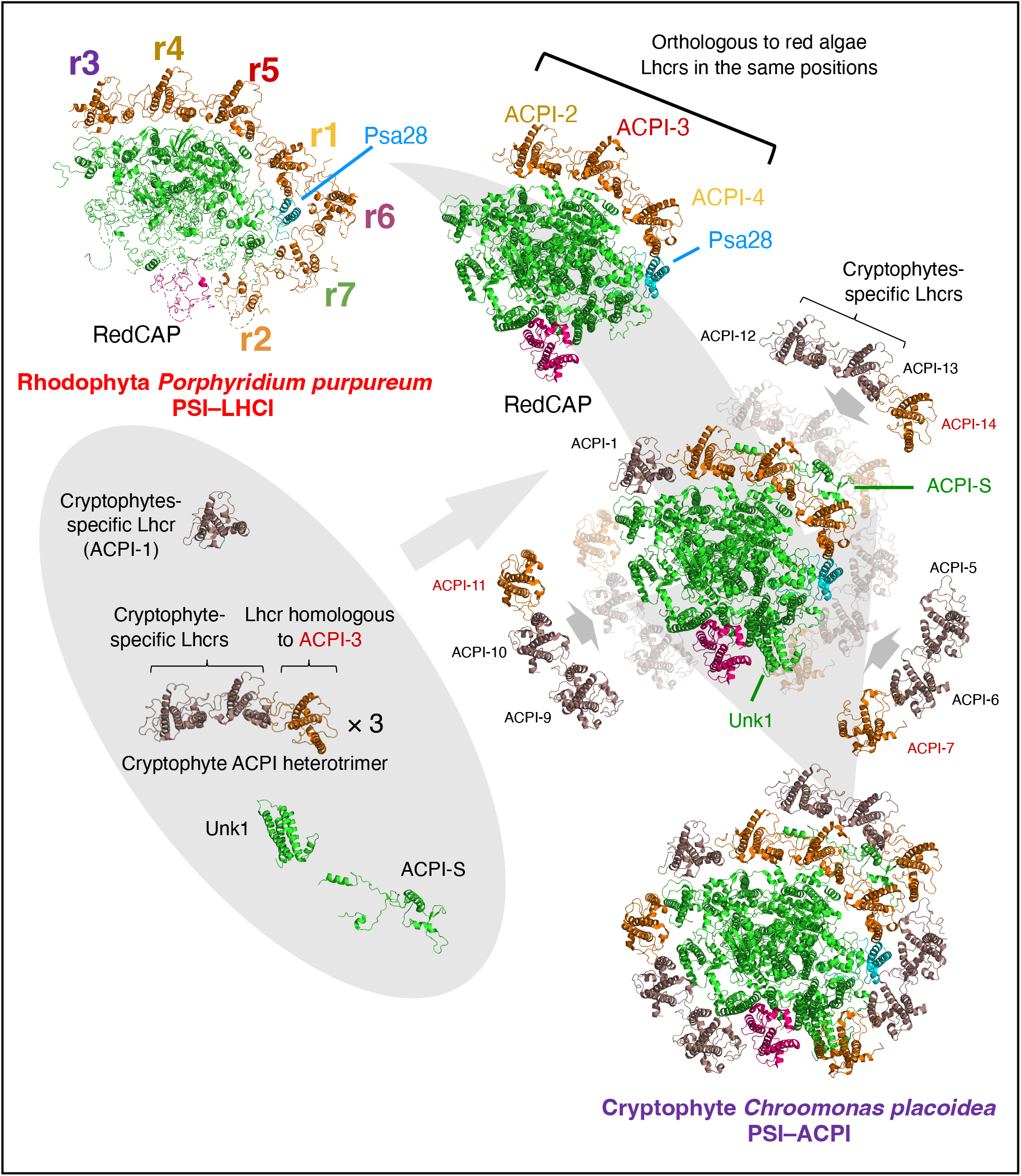
Estimated trajectory of PSI–LHCI evolution of Cryptophytes after endosymbiosis with red alga based on the molecular phylogeny of LHCI. The PSI–ACPI of Cryptophytes *Chroomonas placoidea* has one RedCAP and 13 LHCs, including five Lhcrs homologous to red algal Lhcrs and eight Cryptophyte-specific Lhcrs. Color code is the same as in Figure 5. The cryptophyte PSI–ACPI conserved the PSI core including Psa28 with RedCAP and ACP-4, 3, and 2 at positions p0, p4–p6, respectively, as the putative PSI–LHCI in ancestral red alga. ACP-2–4 are orthologous to PpLhcr1, r5,

Not all Lhcrs in a diatom PSI–FCPI directly descend from red algal LHCI. The diatom *Chaetoceros gracilis* PSI–FCPI possesses either none or one RedCAP molecule and 16 or 23 FCPs including eight Lhcrs (Nagao *et al*., 2020; Xu *et al*., 2020). In the structure of *C. gracilis* PSI–FCPI, positions p0–p7 are occupied by CgRedCAP, and CgLhcr1–r7. CgRedCAP at p0 is homologous to PpRedCAP at p0 of *P. purpureum* PSI–LHCI. CgLhcr1 and CgLhcr5, belonging to groups I and V, respectively, are positioned in p1 and p5, consistent with LHCIs belonging to groups I and V at p1 and p5 in red algal PSI–LHCI. However, all other positions except for p4 are occupied by Lhcrs belonging to Stramenopiles/Haptophyta-specific groups. Furthermore, CgLhcr8 and CgLhcr10, belonging to the Stramenopiles/Haptophyta-specific group I, are positioned counterclockwise adjacent to CgLhcr7 (positions p8 and p9, respectively). The FCPI assigned at p4 in *C. gracilis* PSI–FCPI is CgLhcr4, belonging to the Lhcq subfamily (Nagao *et al*., 2020; Kumazawa *et al*., 2022). Altogether, the diatom PSI– FCPI shares orthologous LHCIs only at p0, p1, and p5 with the red algal PSI–LHCI. This suggests that diatoms have lost many LHCIs from red algae and replaced them with diversified FCPIs during endosymbiosis of red algae.

## Discussion

The molecular assembly model of the red-lineage PSI–LHCI has been discussed only based on spatial arrangements of the subunits in the structures (Bai et al., 2021; Zhao et al., 2023). However, this study made it clear that it is necessary to consider an evolutionary model of the photosynthetic supercomplex using both molecular phylogeny and structural information. Here, we would like to propose new evolutionary trajectory of LHCI proteins associated with PSI in red-lineage algae.

### Putative composition of LHCIs in the common ancestor of primitive red algae

The smaller number of LHCI in *C. merolae* PSI can be due to genome reduction in Cyanidioschyzonales, including *C. merolae*, and Cyanidiales, resulting in small genome sizes and numerous gene deletions (Cho *et al*., 2023). Rhodophytina and Cyanidiophyceae diverged from the common ancestor of red algae. Rhodophytina had a conserved set of Lhcrs: groups I–VII. However, Cyanidiophyceae, Cyanidioschyzonales, and Cyanidiales, have only three LHCIs which belong to groups V–VII. Galdieriales have six Lhcr molecular species belonging to groups I, IV–VII, and the Galdieriales-specific clade. Furthermore, Rhodophytina and Galdieriales possess RedCAP in their genomes, whereas Cyanidioschyzonales and Cyanidiales do not (Engelken *et al*., 2010; Sturm *et al*., 2013). This suggests that only a few Lhcr genes remained after genome reduction in Cyanidioschyzonales and Cyanidiales, and that they would ensure PSI antenna size by repurposing the same genetic products, CmLhcr1 and CmLhcr2, to different position around PSI.

Considering that Galdieriales is the earliest order diverged from others in Cyanidiophyceae, it is reasonable to deduce the LHCI composition of the last common ancestor of red algae from those of Rhodophytina and Galdieriales. Since both Rhodophytina and Galdieriales possess RedCAP, the ancestral Rhodophyta likely had RedCAP at p0. Galdieriales has Lhcrs of group I, IV–VII, which are orthologous Lhcrs in *P. purpureum* PSI– LHCI bound at p1, p4–p7, while it lacks group II and III Lhcrs. Galdieriales conserves the Psa28 (XP_005707993.1, called PsaR, synonymously) subunit, which was suggested to stabilize LHCIs at p2–p4 in *P. purpureum* PSI–LHCI (You *et al*., 2023). In Galdieriales, Psa28 may help the association of at least LHCI at p4. Thus, the ancestral Rhodophyta should have a conserved fundamental Lhcrs–RedCAP composition with RedCAP at p0 and Lhcrs at least at p1, and p4–p7. This also suggests that *C. merolae* PSI–LHCI is not a “primitive” PSI– LHCI.

### Evolutionary path of LHCI from red algae to Stramenopiles/Haptophyte through Cryptophyta

The evolutionary model of Cryptophyta PSI–ACPI introduced in Figure 6 suggest its progression from the ancestral red algae PSI–LHCI. Cryptophyta acquired a plastid from red algae, having its endosymbiotic nucleus, the nucleomorph, derived from red algae (Van Der Auwera *et al*., 1998; Archibald *et al*., 2001). This supports the direct descendance of Cryptophyte PSI–ACPI from red algae PSI–LHCI, without additional endosymbiosis. The ancestral red algae PSI–LHCI, featuring LHCI at p0, 1, and p4–p7, shares positions p0, p4– p5 with Cryptophyte PSI–ACPI. Despite losing group I Lhcr, Cryptophyta has diversified its Lhcrs not only to fill position p7 but also to expand its antenna size by creating three heterotrimers. This research meticulously details the evolution of LHCI from primitive red algae to Cryptophytes, presenting Cryptophytes as a prime example of extensive LHCI rearrangement and antenna enlargement of PSI through endosymbiosis.

In contrast, the evolutionary trajectory of Stramenopiles and Haptophyte LHCI is not as straightforward as that of Cryptophyte ACPIs. When considering the evolutionary model of PSI–LHCI based on molecular phylogeny, Stramenopiles, including diatoms and Haptophytes, acquired the Lhcq subfamily in addition to the Lhcr subfamily (Kumazawa *et al*., 2022). In diatom FCPI, only group I and V Lhcrs (CgLhcr1 and CgLhcr5) at p1 and p5, respectively, have conserved positions from red algae. Other Stramenopiles and at least Pavlovales in Haptophyta share group-I and -V Lhcrs. In diatom PSI–FCPI, Lhcrs at p2, p3, p6–p9 belong to the Stramenopiles/Haptophyta-specific groups(Nagao *et al*., 2020; Kumazawa *et al*., 2022). CgLhcr4, which does not belong to the Lhcr subfamily but to the Lhcq subfamily, is assigned to p4 in diatom PSI–FCPI. Moreover, the Lhcq subfamily in PSI–FCPI not only binds directly to the PSI core but also forms the outer layer of LHCs in PSI–FCPI, contributing to a larger antenna size of the diatom PSI (Nagao *et al*., 2020; Xu *et al*., 2020; Kumazawa *et al*., 2022). This indicates that Stramenopiles has experienced a distinct process of LHCI re-acquisition around PSI, which would be largely different from that of Cryptophytes.

The evolutionary history from red algae to red-lineage algae has been previously described (Stiller *et al*., 2014). Linear regression analysis on nuclear genomes suggested serial endosymbiosis in red-lineage algae: Cryptophyta engulfed a red alga as secondary (2nd) endosymbiosis; the photosynthetic Stramenopiles incorporated Cryptophyta as tertiary (3rd) endosymbiosis; Haptophyta acquired many genes from Stramenopiles as quaternary (4th) endosymbiosis. However, the plastid-encoded genes of Haptophyta and Cryptophyta strongly support monophyly in the phylogenetic tree (Kim *et al*., 2017). Haptophyta may have acquired the ancestral Cryptophyta plastid before or after massive gene transfer from Stramenopiles, reconciling the two seemingly contradictory phylogenetic trees of different genomes (Dorrell *et al*., 2017; Dorrell *et al*., 2021; Penot *et al*., 2022). LHC genes are nuclear-encoded and should follow the history of nuclear-encoded genes derived from the quaternary endosymbiosis (Stiller *et al*., 2014; Dorrell *et al*., 2017). Consistently, the composition of LHC subfamilies of Haptophyta is similar to that of Stramenopiles (Kumazawa *et al*., 2022). In contrast, most genes coding for the PS core are encoded in the plastid genome originated from Cryptophyta. Based on these facts, we hypothesize that the origin of the genes coding for the Haptophyte PS core complex and that for light-harvesting antennas can be chimeric. Further genetic and structural analysis of the PSI–LHCI in Haptophytes is required to confirm/dismiss this hypothesis.

### Molecular Evolution of PS supercomplexes through “neolocalization”

Gene duplication and functional diversification, for instance in biosynthetic enzyme families, are typically referred to as “neofunctionalization” (e.g. Hansen et al. 2021). However, even after intensive diversification, the primary function of the LHC family and RedCAP remains light-harvesting. Therefore, their diversification can be best considered as that of structural relocalization rather than function. In our study on the red alga Cyanidiales *Cyanidium caldarium* PSI–LHCI (Kato *et al*., 2024), we proposed the term “neolocalization” (Kato *et al*., 2024). It was defined as a phenomenon where a structural defect caused by gene loss is complemented or modified by the product of another existing gene. In this study, we expand the definition of neolocalization to include modifications by the product of a gene differentiated after duplication from an existing one. Phenomena matching neolocalization are also observed in the green lineage. In green algae *Chlamydomonas reinhardtii*, green algal Lhca7 and terrestrial plant Lhca2 do not share orthology, yet they bind at the same position in green-lineage PSI–LHCIs (Neilson and Durnford, 2010; Suga *et al*., 2019; Ben-Shem *et al*., 2003).

For the PS supercomplex, which has a long evolutionary timeframe, drastic events like genome evolution and secondary endosymbiosis may primarily trigger neolocalization, driving the molecular evolution of LHCs. To understand this process, a general model of molecular evolution must consider both molecular phylogeny and structure. At present, the complete evolutionary paths of LHC diversification and differentiation from Rhodophytina red algae to Stramenopiles and Haptophyta through Cryptophyta remain to be elucidated. As more genomic information and more structural models become available, the relationship between these taxa will become clearer, allowing the construction of an entire evolutionary model of PS supercomplexes.

## Methods

### LHC Protein Sequence Acquisition

LHC protein sequences were collected from genomes or transcriptomes of diverse red-lineage species, including 37 Rhodophyta comprising 33 species, five diatoms, two Eustigmatophyceae, four Haptophytes including one Pavlovophyceae, three Phaeophyceae (Brown algae) and three Cryptophytes. Thirty-three species of red algae include one Cyanidiales, two Cyanidioschyzonales, six Galdieriales, two Rhodellophyceae, two Compsopogonophyceae, two Stylonematophyceae, two Porphyridiophyceae, two Bangiophyceae, and 18 Florideophyceae. These genomic and transcriptomic datasets were accessed from databases such as ChaetoBase v1.1, NCBI RefSeq, NCBI GenBank, NCBI SRA, PDB, JGI Phycocosm and 1KP (https://db.cngb.org/onekp/, https://ftp.cngb.org/pub/gigadb/pub/10.5524/100001_101000/100627/assemblies/). Specific details of species and corresponding references are provided in Supplemental Table I. For most diatoms, LHCs had been previously annotated, except for *Fistulifera solaris* JPCC DA0580 (Kumazawa *et al*., 2022). The protein sequence of *Thalassiosira pseudonana* Lhcr18 was modified as the amino-acid sequence of g8189.t1 in the updated *T. pseudonana* genome (https://doi.org/10.5683/SP2/ZDZQFE) (Filloramo *et al*., 2021). A BLAST similarity search was used to procure LHCs from various lineages, adapting techniques from Kumazawa et al. (2022). Then, the reference sequences used in the BLASTP search were replaced with the Lhcrs in *Porphyridium purpureum*. The transcriptomes of 1KP included 28 species of red algae, among which 23 species were selected for the next analyses based on the quality of the LHC alignment (Leebens-Mack *et al*., 2019). Contaminated sequences in the 1KP dataset were identified and eliminated using molecular phylogenetic analysis and BLASTP searches against the NCBI nr database. The IsoSeq transcriptomes of *Cyanidiococcus yangmingshanensis* 8.1.23 F7 and *Cyanidium caldarium* DBV 063 E5 were obtained from NCBI SRA and translated using TransDecoder (https://github.com/TransDecoder/TransDecoder) and LHC proteins were identified using BLASTP and clustered manually based on MAFFT alignment (Cho *et al*., 2023; Katoh and Standley, 2013). LHCs belonging to the Lhcr subfamily of secondary endosymbiotic algae were obtained by preliminary phylogenetic analysis using muscle5 with super5 mode for alignment, ClipKit v1.4.1 with kpic-smart-gap mode for trimming, and IQ-TREE v2.2.2.7 to infer a phylogenetic tree (Edgar, 2022; Steenwyk, Buida, Li, X.,-X., Shen, *et al*., 2020; Minh *et al*., 2020). All sequences for the following analyses were carefully curated for LHC conserved domain; some were modified at their N-terminal region and C-terminal region. All modified sequences are explicitly indicated in the figure 2 and 4.

Lhca sequences of *Prasinoderma coloniale* belonging to the earliest divergent taxa of green algae were obtained for the root in the phylogenetic tree (https://ftp.cngb.org/pub/CNSA/data2/CNP0000924/CNS0223647/CNA0013964/) (Li *et al*.,

2020).

### Phylogenetic Analysis of LHC

LHC sequences were aligned using MAFFT E-INS-I v7.490 (Katoh and Standley, 2013), which is optimized for multi-domain proteins. Sequence alignments were then refined using ClipKit v1.4.1 (Steenwyk, Buida, Li, X., X., Shen, *et al*., 2020) with kpic-smart-gap mode for the following tree inference. The molecular phylogenetic trees were inferred using IQ-TREE v2.2.2.7 with extended model selection (-m MFP option) (Minh *et al*., 2020; Kalyaanamoorthy *et al*., 2017). For exhaustive tree topological search, 500 initial parsimony trees were constructed, the number of tree search iterations was extended to 1,000, and perturbation strength was specified to 0.7 as IQ-TREE parameters. The inferred trees were visualized using iTOL v6 (Letunic and Bork, 2021).

### Visualization of PSI–LHCI structure

All models of PSI ‒ LHCI structures were obtained from RCSB PDB (https://www.rcsb.org/). The following models were acquired: a *Cyanidioschyzon merolae* PSI–LHCI (ID: 5ZGB), a diatom *Chaetoceros gracilis* PSI–FCPI (ID: 6LY5), a red alga *Porphyridium purpureum* single-PBS-PSII-PSI-LHCs megacomplex (ID: 7Y5E), and a Cryptophyte *Chroomonas placoidea* PSI–ACPI (ID: 7Y7B). The models of the complexes were visualized using Open-Source PyMOL v2.5.0 (Schrodinger LLC, 2015).

## Acknowledgments

This work was supported in part by JSPS KAKENHI grant Nos. JP22KJ2017 (M.K.), JP23H02347 (K.I.), and a grant from Institute for Fermentation, Osaka, Japan (K.I.). We would like to thank Dr. Atsushi Takabayashi of Hokkaido Univ. and Dr. Ryo Nagao of Shizuoka Univ. for their valuable discussions throughout this study.

## Author Contributions

M.K. and K.I. conceived the project; M.K. performed all analyses; M.K. drafted the original manuscript; M.K. and K.I. revised the manuscript and wrote the final manuscript, and both authors joined the discussion of the results.

## Additional information

Supplemental Table 1: Accessions of genomic or transcriptomic data.

Supplemental Data 1: LHC protein sequences used in the analyses.

Supplemental Data 2: Alignment to infer the phylogenetic tree of Figure 3.

Supplemental Data 3: Alignment to infer the phylogenetic tree of Figure 5.

Supplemental Data 4: Newick file of the phylogenetic tree of Figure 3.

Supplemental Data 5: Newick file of the phylogenetic tree of Figure 5.

